# A genome-wide resource for high-throughput genomic tagging of yeast ORFs

**DOI:** 10.1101/226811

**Authors:** Matthias Meurer, Yuanqiang Duan, Ehud Sass, Ilia Kats, Konrad Herbst, Benjamin C Buchmuller, Verena Dederer, Florian Huber, Daniel Kirrmaier, Martin Štefl, Koen Van Laer, Tobias P Dick, Marius K Lemberg, Anton Khmelinskii, Emmanuel D Levy, Michael Knop

## Abstract

Here we describe a C-SWAT library for high-throughput tagging of *Saccharomyces cerevisiae* ORFs. It consists of 5661 strains with an acceptor module inserted after each ORF, which can be efficiently replaced with tags or regulatory elements. We validate the library with targeted sequencing and demonstrate its use by tagging the yeast proteome with bright fluorescent proteins, determining how sequences downstream of ORFs influence protein expression and localizing previously undetected proteins.

---

Genome-wide libraries of strains where every open reading frame (ORF) is fused to a constant tag are valuable resources for proteome-wide studies in *Saccharomyces cerevisiae*. Different libraries are available to assess properties such as protein localization, abundance, turnover and protein-protein interactions for a large fraction of the yeast proteome^1-6^. However, construction of such libraries is costly and time-consuming, which hampers genome-wide endeavors with improved tags such as novel fluorescent proteins, tags bearing sequences for RNA detection^7^ or regulation of gene expression^8^.

To overcome these limitations, we recently developed the SWAp-Tag (SWAT) approach for high-throughput tagging of yeast ORFs and used it to N-terminally tag proteins of the endomembrane system^9^. This approach requires a one-time construction of SWAT strains where individual ORFs are marked with an acceptor module (Fig. 1a). New strains can be rapidly derived from SWAT strains using automated procedures to replace the acceptor module with practically any tag or regulatory element provided on a donor plasmid^9^ (Fig. 1a).

**Figure 1.**
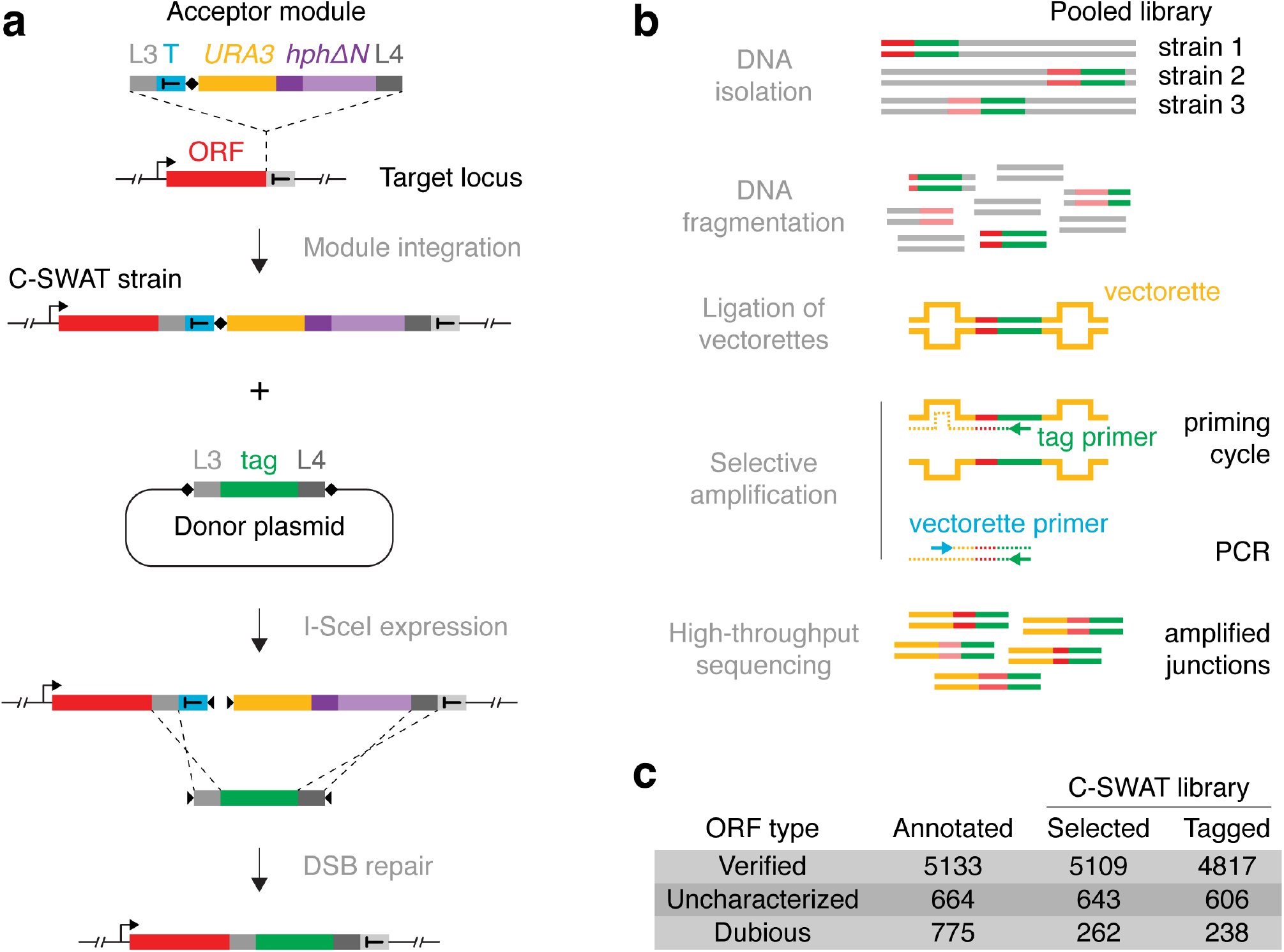
Design, construction and validation of the C-SWAT library. **(a)** Outline of the C-SWAT approach. A C-SWAT acceptor module is inserted into the genome before the stop codon of yeast ORFs. A construct for conditional expression of the I-SceI endonuclease (not shown) and a donor plasmid carrying the desired tag are then introduced into the C-SWAT strains by transformation or genetic crossing. Upon expression, I-SceI induces double strand breaks (DSBs) at the indicated positions (♦) in the acceptor module and the donor plasmid. DSB repair by homologous recombination leads to replacement of the acceptor module by the tag. **(b)** Outline of the Anchor-Seq targeted sequencing approach. A library of strains with different ORFs (red) modified with a constant tag (green, *e.g*. C-SWAT acceptor module) is pooled. Vectorette adaptors are ligated to sheared genomic DNA (Fig. S1a). A primer annealing to the tag sequence initiates a first cycle of DNA synthesis. Synthesis extends into the adaptor, which creates an antisense strand complementary to the vectorette primer (Fig. S1b) and allows selective amplification of ORF-tag junctions. **(c)** Composition of the C-SWAT library. ORFs are classified as *verified, uncharacterized* or *dubious* (unlikely to encode a functional protein) according to the July 2016 annotation of the *S. cerevisiae genome*.

To apply the SWAT approach to the whole yeast proteome, here we introduce a genome-wide C-SWAT library. This library enables high-throughput genome engineering at 3’ ends of yeast ORFs and can be used for high-throughput C-terminal protein tagging. We constructed C-SWAT strains using conventional PCR targeting^10,11^ to insert a C-SWAT acceptor module before the stop codon of individual ORFs at endogenous chromosomal loci (Fig. 1a). The acceptor module consists of homology arms (L3 and L4, for subsequent recombination with the desired tag), a heterologous transcription terminator (T), a restriction site for the I-Scel endonuclease (♦), a selection/counter-selection marker (*URA3*) and a second truncated selection marker (*hphΔN*) (Fig. 1a, Supplementary Information).

To verify correct integration of the acceptor module in each C-SWAT strain, we developed a high-throughput targeted sequencing approach (Anchor-Seq) to sequence the junctions between the 3’ end of each tagged ORF and the 5’ end of the acceptor module (Fig. 1b). In Anchor-Seq, genomic DNA is isolated from a pooled library of strains, where a different ORF is modified in each strain. The junctions of interest are then selectively amplified using vectorette PCR^12^ and subjected to high-throughput sequencing (Fig. 1b, S1, Supplementary Information). We performed Anchor-Seq on pools of six replicates of the C-SWAT library, corresponding to six independent transformants for each ORF. In total, we obtained validated C-SWAT strains for 94% of *verified* or *uncharacterized S. cerevisiae* ORFs and for 238 *dubious* ORFs (Fig. 1c, Table S1).

To tag ORFs with the C-SWAT library, a construct for conditional expression of the I-SceI endonuclease and a donor plasmid carrying the desired tag can be introduced into C-SWAT strains in high-throughput by automated genetic crossing with a donor strain^9^ (Fig. 1a, Supplementary Information). Three types of donor plasmids with different selection strategies can be used: type I for seamless replacement of the acceptor module with the tag and counter-selection for the loss of the acceptor module, type II for selection of tagging events via reconstitution of the hygromycin resistance marker (*hph*) and type III, which introduces the tag together with a new selection marker (Fig. S2a). We estimated the tagging efficiency with these strategies using C-SWAT strains for 20 highly expressed genes. We observed an average tagging efficiency of ~98% with a type I donor and > 99% with the other two donor types (Fig. S2b). This demonstrates that the C-SWAT library can be used for high-throughput strain construction without the need for subsequent clonal selection. We note that for 1-4% of ORFs, endogenous repetitive sequences surrounding the tag integration site could interfere with seamless tagging using the C-SWAT approach^13^.

The yeast GFP library^1^, in which 4159 ORFs are tagged with GFP(S65T)^14^, has been widely used to study the yeast proteome. However, since the construction of this library, various fluorescent proteins with improved properties have been developed. The C-SWAT library provides a platform to profit from these developments. We found that in yeast the green fluorescent protein mNeonGreen^15^ and the red fluorescent protein mScarlet-I^16^ are up to three times brighter than fluorescent proteins used in previous libraries^1,4,9^ (Fig. S3). Using the C-SWAT library, we tagged the yeast proteome with mNeonGreen and mScarlet-I, generating three new libraries: mNG-I (seamless tagging with mNeonGreen), mNG-II and mSC-II (where mNeonGreen and mScarlet-I are followed by a heterologous terminator) (Fig. 2a). We determined the expression levels of proteins tagged in these libraries using fluorescence measurements of colonies^4^. Over 4300 proteins were expressed at detectable levels (> 1.2 fold above background) in each library (Fig. 2b, Table S2). This is consistent with the number of proteins detected with mass spectrometry in yeast grown under standard laboratory conditions^17^. Protein expression levels correlated well between the three libraries (Fig. S4a) and with independent estimates of protein abundance^17^ (Fig. S4b), demonstrating the reproducible and reliable nature of proteome-wide tagging with the C-SWAT library.

**Figure 2.**
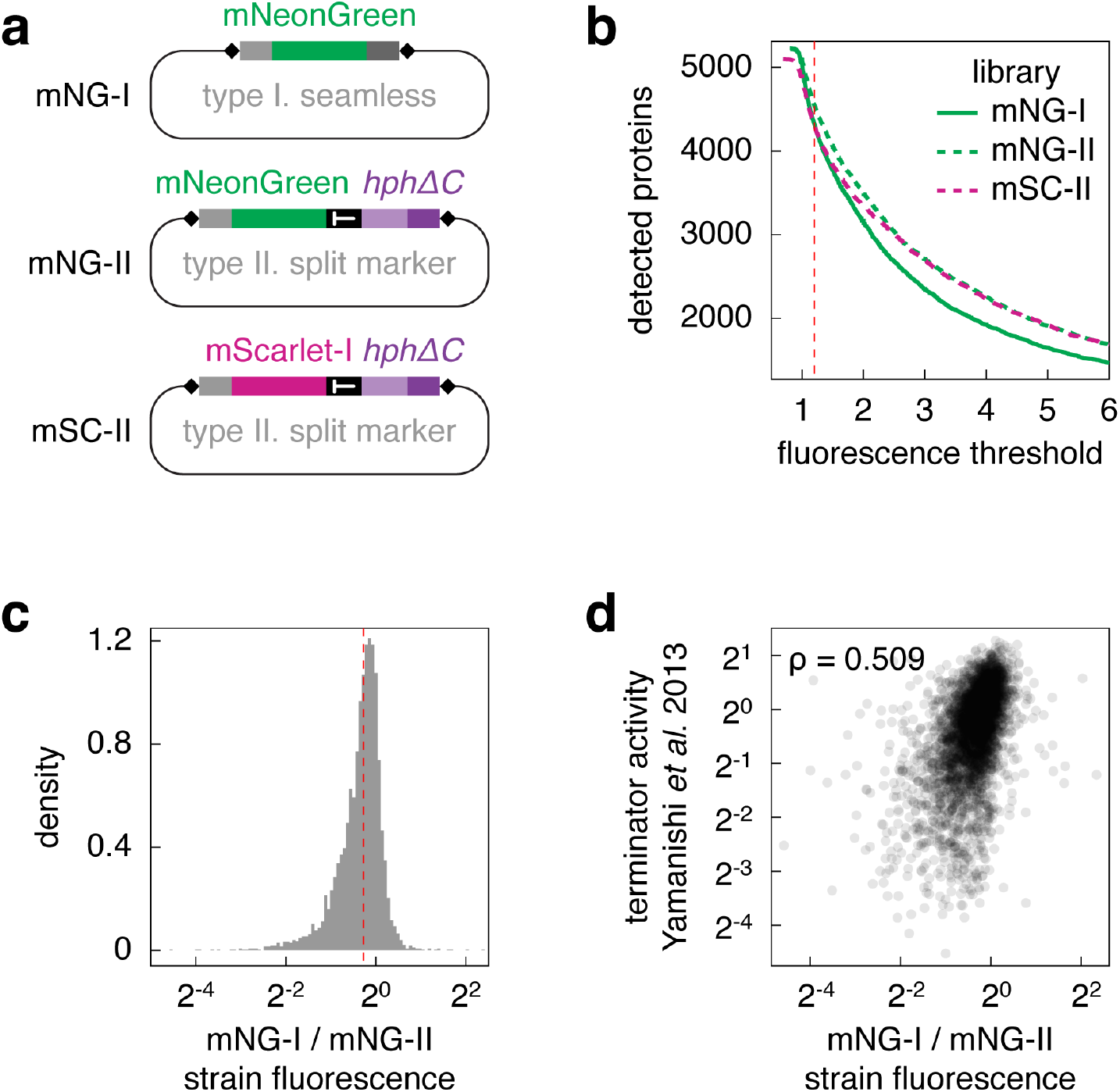
High-throughput protein tagging with the C-SWAT library. **(a)** Donor plasmids used to tag the yeast proteome with mNeonGreen and mScarlet-I fluorescent proteins using the C-SWAT library. **(b)** Number of protein fusions detected in each library with fluorescence measurements of colonies. 4312, 4537 and 4301 strains in the mNG-I, mNG-II and mSC-II libraries, respectively, had a fluorescence signal at least 1.2 fold above background (red dashed line). **(c)** Distribution of differences in protein expression levels (strain fluorescence corrected for background) between mNG-I and mNG-II libraries. Median, 0.83 (red dashed line). **(d)** Correlation of endogenous transcription terminator activity for each ORF, measured in ref. 18, and differences in protein expression levels between mNG-I and mNG-II strains.

Having libraries with seamless and non-seamless protein tags allowed us to examine how regulatory elements downstream of each ORF contribute to protein expression. We observed that protein levels were on average ~20% higher in mNG-II strains (non-seamless tagging) compared to mNG-I strains (seamless tagging) (Fig. 2c). Protein levels differed by more than two fold for ~11% of the proteome, with 466 and 10 proteins exhibiting lower and higher expression in mNG-I strains, respectively. Moreover, the difference between mNG-I and mNG-II strains correlated with the strength of the endogenous transcription terminator for each ORF^18^ (Spearman’s rank correlation coefficient = 0.509, Fig. 2d). Consistent with these observations, the heterologous *ADH1* terminator used in mNG-II and mSC-II libraries is stronger than terminator sequences of most yeast ORFs^18^. Together these results demonstrate that tagging modules with heterologous terminators, commonly used for C-terminal protein tagging in yeast^19,20^, can measurably impact protein expression and suggest applications of the C-SWAT library to study regulation of gene expression.

We observed expression of 208 proteins in the mNG-I and mNG-II libraries that were previously undetected with various independent approaches^17^ (Fig. 3a). Compared to the entire C-SWAT library (Fig. 1c), this group is enriched in ORFs annotated as *uncharacterized* (62 ORFs) or *dubious (i.e*., unlikely to encode functional proteins based on available data, 133 ORFs). We used fluorescence microscopy to examine the localization of 60 such proteins. Notably, for 9 of them (5 *uncharacterized* and 4 *dubious* ORFs) we could detect expression and a specific non-cytosolic localization even in mNG-I strains, where transcription is not influenced by a heterologous terminator (Fig. 3b, c, Table S2), suggesting that these are indeed functional proteins. With updates to the yeast genome annotation, 80 of 238 *dubious* ORFs in the C-SWAT library were recently reclassified as *verified* or *uncharacterized* (Table S2). We note that the reclassified and the remaining *dubious* ORFs exhibit similar expression levels when tagged with mNeonGreen (Fig. S4c), raising the possibility that more *dubious* ORFs actually encode functional proteins^21^.

**Figure 3.**
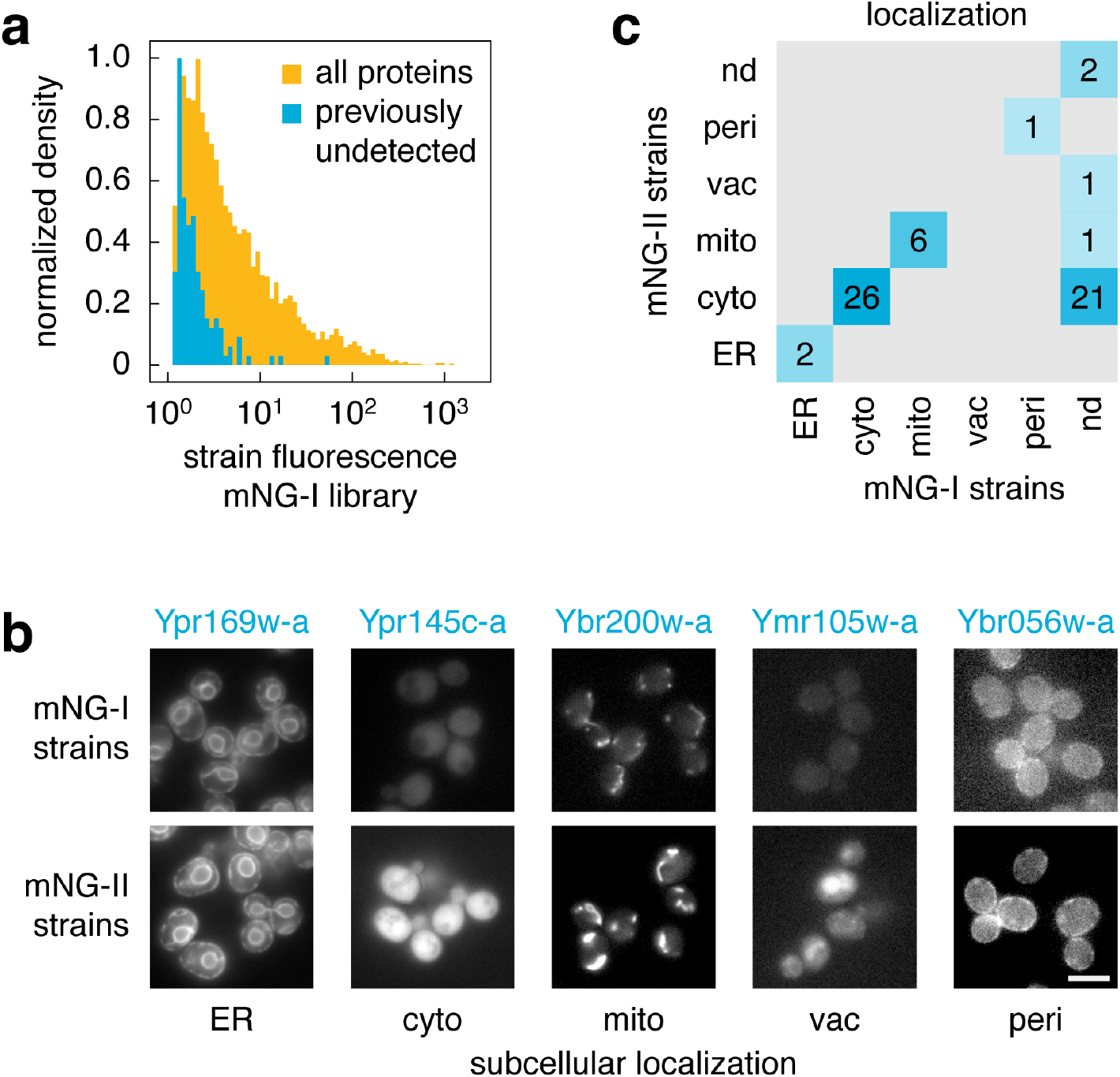
Localization of previously undetected proteins. **(a)** Fluorescence levels of mNG-I strains (fold over background) expressing previously undetected proteins. Fluorescence measurements of the entire mNG-I library are shown for comparison. Only strains with fluorescence at least 1.2 fold above background were considered. **(b, c)** Fluorescence microscopy of strains from mNG-I and mNG-II libraries expressing 60 previously undetected proteins tagged with mNeonGreen. (b) Examples of fusions with different subcellular localizations. The two images for each protein were acquired and processed identically. Scale bar, 5 μm. (c) Summary of observed subcellular localizations. ER (endoplasmic reticulum), cyto (cytosol), mito (mitochondria), vac (vacuole), peri (cell periphery) and nd (expression not detected).

In conclusion, the C-SWAT library is a versatile resource for exploring the yeast genome and proteome. With this tool at hand, the ORFeome can be efficiently manipulated to generate libraries with a variety of tags for protein or RNA detection, to study regulation of gene expression or to explore genomic position effects. It is our hope that the simplicity and cost-effectiveness of C-SWAT will make construction of custom genome-wide libraries routine and facilitate systematic studies.

## Supplementary Information

Supplementary Information contains Methods, Supplementary Figures and Supplementary Tables.

## Acknowledgments

This work was supported by the Deutsche Forschungsgemeinschaft (DFG) Collaborative Research Center SFB1036 (MK, AK, TPD, MKL), the Weizmann Institute of Science (ES and EDL), the China Scholarship Council (YD), fellowships from the HBIGS graduate school (IK, KH and FH), an SFB1036 travel grant (ES), the Alexander von Humboldt Foundation (MŠ) and, partially, by the DFG grant KN498/11-1 (MK), the I-CORE Program of the Planning and Budgeting Committee grants 1775/12 and 2179/14 (EDL) and the HFSP Career Development Award CDA00077/2015 (EDL). We thank the CellNetworks Deep Sequencing Core Facility (Heidelberg University), the Genomics Core Facility (EMBL) and acknowledge generous support of the DFG for data storage (LDSF2).

## Author contributions

MK, MM and AK planned the work. YD and MM, together with ES, constructed the library, with help from BCB, VD, KH, FH, DK, IK, MŠ, KVL, AK and MK. ES and EDL developed Anchor-Seq and EDL analyzed the sequencing data, with input from MM and KH. YD, MM, IK, AK and DK generated and analyzed the mNeonGreen and mScarlet-I libraries. AK and MK wrote the manuscript with EDL and ES, with input from all authors.

## Supplementary Information

### Methods

#### 1. Construction of the C-SWAT library

##### Acceptor module

The acceptor module used to construct the C-SWAT library (plasmid pMaM471, Table S3) is composed of the following elements:

- linker L3 (5′-<u>cgtacgctgcaggtcg</u>acggtggcggttctggcggtggcggatcc-3′), which contains the <u>S3 primer annealing site</u> for gene tagging by PCR targeting;
- STOP codon;
- terminator sequence of the *CYC1* gene from *Saccharomyces paradoxus;*
- the recognition sequence for the I-SceI endonuclease;
- the *URA3* gene with its endogenous promoter and terminator from *Saccharomyces cerevisiae*;
- *hphΔN* sequence coding for a C-terminal fragment (amino acids 146-342) of the *hph* (hygromycin resistance-encoding gene) marker;
- terminator sequence of the *ALG9* gene from *Saccharomyces paradoxus;*
- linker L4 (5’-agttcttctttgagatatcgattgaa<u>cgagctcgaattcatcgat</u>-3’), which contains the <u>S2 primer annealing site</u> for gene tagging by PCR targeting.

##### Library background strain

The strain BY4741 (ref. 1, Table S4), used to construct the most popular collections of yeast strains such as the knockout and GFP libraries^2,3^, was chosen as the background strain for the C-SWAT library. All additional elements required for the SWAT procedure (a donor plasmid and a construct for inducible expression of the I-SceI endonuclease) can be introduced into C-SWAT acceptor strains by direct transformation or by genetic crossing with the donor strain YMaM639 (Table S4) carrying the desired donor plasmid.

##### Donor strain and plasmids

The donor strain YMaM639 was constructed using the strain Y8205 (ref. 4, Table S4). In this strain the *leu2Δ0* locus carries the *GAL1pr-NLS-I-SceI-natNT2* construct for galactose-inducible expression of the I-SceI endonuclease with a nuclear localization signal (NLS), which increases the efficiency of tag swapping. In addition, this strain also contains the *can1Δ::STE2pr-SpHIS5* and *lyp1Δ::STE3pr-LEU2* markers for selection of *MAT***a** or *MAT***alpha** haploids at the end of the automated genetic crossing procedure using synthetic genetic array (SGA) methodology.

Three different template plasmids were used to construct donor plasmids (Table S3):

- pMaM482 – type I donor template (for seamless tag swap);
- pMaM484 – type II donor template (for tag swap with reconstitution of the *hph* marker);
- pMaM496 – type III donor template (for tag swap with introduction of the *kanMX6* marker).

The backbone of donors type I and II is pRS41K (ref. 5) and pRS41 (ref. 6) for donor type III. Tags can be cloned into these templates *e.g*. via *Bam*HI + SpeI restriction enzyme cut sites. Here we constructed three donors for tagging with the mNeonGreen fluorescent protein (type I/II/III: pYD10/11/14) and one donor for tagging with the mScarlet-I fluorescent protein (pYD13, type II donor) (Table S3). All plasmids and sequences are available upon request.

##### Selection of ORFs and primer design

We selected 6071 yeast open reading frames (ORFs) for tagging with the acceptor module: all 5797 verified or uncharacterized ORFs and 274 dubious ORFs that do not overlap with any verified or uncharacterized ORFs (retrieved from the *Saccharomyces* Genome Database on July 2016). ORFs from the mitochondrial genome and the 2μ plasmid were not included.

For each ORF, S2/S3 primers for PCR amplification of the tagging module were designed as previously described^7^. For 4081 ORFs, primers synthesized based on the yeast genome sequence from December 2009 (*Saccharomyces* Genome Database) were available from a previous study^8^. For the remaining 1990 ORFs, primers were designed based on the yeast genome sequence from July 2016. In this second set, ORFs with identical S2 and S3 primer sequences were identified (*e.g., HXT15* and *HXT16*). For such cases, only one set of primers was synthesized and assigned to the first ORF by alphabetical order of systematic names. This reduced the set of 1990 ORFs to 1933. The 6014 pairs of S2/S3 primers (Table S1) were obtained from IDT (Integrated DNA Technologies) in 96-well format, such that each well contained a mixture of S2/S3 primers for a different ORF at 5 μM concentration.

##### Strain construction

The acceptor module was amplified by PCR in 96-well format using plasmid pMaM471 as template and ORF-specific S2/S3 primers in each well, as follows. Cooled 96-well PCR plates (4titude, 4ti-0960) were filled with 40 μl/well of a PCR mix using a Liquidator 96 channel manual pipettor (Mettler Toledo):

- 5 μl of 10x HiFi-buffer (200 mM Tris-HCl, pH 8.8; 100 mM (NH4)2SO4; 100 mM KCl; 1% (v/v) Triton X-100; 1 mg/ml BSA);
- 0.5 μl of 100 mM stock of dNTPs (Bioline, BIO-39026);
- 0.15 μl of 1 M stock MgCl_2_;
- 5 μl of 5 M stock of betaine (Sigma-Aldrich, 61962);
- 0.5 μl of template DNA (200 ng/μl stock);
- 27.85 μl of H2O;
- 1 μl of a high fidelity DNA polymerase (self-made)).

A mixture of ORF-specific S2/S3 primers (10 μl of 5 μM stock) was added to each well from 96-well primer source plates using the 96 channel manual pipettor. The plates were sealed with aluminum seals (Steinbrenner Laborsysteme, SL-AM0550). PCR was then carried out in a Biometra TAdvance or TProfessional (Analytik Jena) using the following program:

- 2 min at 95°C;
- 30 cycles of 20 s at 95°C/30 s at 66°C/2 min 30 s at 72°C;
- 5 min at 72°C;
- incubation at 4°C.

Frozen BY4741 competent yeast cells were prepared and transformed with PCR-amplified acceptor modules in 96-well plates as previously described^7,8^. Transformation mixtures were manually plated onto 9 cm petri dishes with SC-Ura agar medium (synthetic complete medium lacking uracil and with 2% (w/v) glucose as carbon source). After 2-3 days of incubation at 30°C, six clones from each transformation were manually streaked for single colonies. Single colonies were then grown in 96-well format in SC-Ura medium with 15% (v/v) of glycerol. The resulting six replicates of the C-SWAT library, one for each clone, were stored at −80°C.

#### 2. Library validation by targeted sequencing (Anchor-Seq)

We developed and used Anchor-Seq to verify correct integration of the acceptor module in each C-SWAT strain. The validation procedure involved pooling of each replicate of the C-SWAT library, extraction of genomic DNA from each pool, DNA fragmentation and size selection, ligation of vectorette adaptors, selective amplification of junctions between the C-SWAT acceptor module and upstream genomic sequences, and Illumina sequencing, as detailed below.

Each of the six C-SWAT library replicates was grown to saturation in 384-well plates (50 μl of YPD medium per well). For each replicate, all strains were pooled, cells were harvested by centrifugation, washed once and re-suspended in 10 ml ddH_2_O, aliquoted and stored at −80°C before further processing.

Genomic DNA was extracted from each pool (300 μl sample) with YeaStar Genomic DNA extraction kit (Zymo-Research, #D2002) according to manufacturer’s protocol using chloroform. The yield and quality of extracted DNA (typically 7-10 μg) were assessed by absorbance at 260 and 280 nm wavelengths, measured with a NanoDrop ND-1000 spectrophotometer. Genomic DNA (5 μg in 125 μl of 10 mM Tris-HCl, pH 8.0) was fragmented in microTUBEs (Covaris, #520045) using a focused ultrasonicator (Covaris, E220x) with shearing parameters set for a fragment size of 800 bp (shearing time of 50 sec per tube, peak incident power of 105 watts, duty factor of 5% and 200 cycles per burst).

Vectorette adaptors were then ligated to the sheared genomic DNA as follows. Single-stranded adaptor oligonucleotides (356-vectorette and 355-vectorette, Table S5) were individually resuspended in annealing buffer (100 mM potassium acetate, 30 mM HEPES-KOH pH 7.5) to 100 μM, mixed in equal amounts in a 1.5 ml Eppendorf tube and annealed (5 min incubation at 98°C, cooling to ~23°C in a water bath over a period of 3 h). Annealed vectorette adaptors were then ligated to sheared genomic DNA (1 μg) using NEBNext Ultra™ II DNA Library Prep Kit for Illumina (New England Biolabs, #E7645S). Ends preparation and A-T ligation were performed according to manufacturer’s protocol using 15 μM of vectorette adaptors. Fragment sizes of 300-800 bp were selected by agarose gel electrophoresis (3.5% Nusieve 3:1 agarose, Lonza #50090) and extracted with Qiaquick gel extraction kit (Qiagen #28704) in 25 μl of 10mM Tris-HCl solution pH 8.

DNA fragments containing the SWAT acceptor module were selectively amplified as follows. First, 20 μl of purified adaptor-ligated fragments were used as input for 15 cycles of PCR with primers 503-Tag_primer and 357-VC_primer (Table S5). The PCR product was purified with Qiaquick PCR purification kit (Qiagen #28104) and used as input for 15 cycles of PCR with primers 382-P5 and 529-BC_P7 (Table S5). This set of primers added the P5 and P7 Illumina sequences and a 6 nucleotide barcode for multiplexed sequencing. Fragment sizes were analyzed on a TapeStation 2200 (Agilent HS D1000 ScreenTape, #5067-5584) and typically followed a normal distribution peaking at 550-650 bp. In case fragments outside of 300-800 bp were present, samples underwent further size selection by agarose gel electrophoresis and gel extraction before Illumina sequencing. The outcome of selective amplification was controlled before sequencing, using quantitative PCR (qPCR) to compare the abundance of fragments corresponding to ORFs that were tagged in the C-SWAT library (*ERG1, ERG11* and *ERV25;* amplification with an ORF-specific primer and a reverse primer in the acceptor module (Tag-rev), Table S5 – qPCR on-target) and ORFs that were not tagged (*ACO1, RPL30* and *ASC1;* amplification with two ORF-specific primers, Table S5 – qPCR off-target). 5 ng of each sample (before and after selective amplification) were assayed using fast SYBR Green master mix (Thermo Fischer Scientific #4385612). The abundance (cycle threshold (CT) values) of on-target and off-target sequences, before and after selective amplification, were determined by StepOnePlus Software v2.3 (Thermo Fischer Scientific). Enrichment was defined as the difference of CT values (ΔCT) and calculated as 2^(***A+B***), where ***A*** is the ΔCT between off- and on-target sequences before selective amplification, and ***B*** is the ΔΔCT for these sequences after selective amplification. We observed enrichments typically in the ~10^5^-10^6^ fold range.

Samples were normalized to 10 nM final concentration and subjected to sequencing (Illumina MiSeq PE300_V3 flowcell). All six libraries were sequenced simultaneously. For demultiplexing, the barcodes added during the second PCR step (using primers 382-P5 and 529-BC_P7) were used. In order to infer which genes were tagged correctly or incorrectly, we constructed expected reads using the 140 bp upstream of the stop codon for each ORF. Sequencing reads were then searched for matches to expected reads using custom Perl scripts. Given an ORF *i*, the search detected strict matches as well as six possible frameshifts (−3, −2, −1, +1, +2, +3) and we denote the corresponding counts as M_i_, F^-3^_i_, F^-2^_i_, F^-1^_i_, F^+1^_i_, F^+2^_i_, F^+3^_i_. We also recorded cases where a strict match was found only for the first (1-70 bp upstream of the stop codon) or second half (71-140 bp upstream of the stop codon) of the sequence, presumably due to mutations that occurred in the half not matched. The corresponding counts are X^1^_i_ (first half) and X^2^_i_ (second half). We used three criteria to identify correct clones:

1. –the number of strict matches had to be above 5;
2. –the number of strict matches had to exceed the total number of erroneous reads;
3. –the number of partial matches could not exceed strict matches by more than 5-fold.

That is, ORF *i* was considered correctly tagged if M_i_ > 5, M_i_ > (F^-3^_i_ + F^-2^_i_ + F^-1^_i_ + F^+1^_i_ + F^+2^_i_ + F^+3^_i_), 5*M_i_ > X^1^_i_ and 5*M_i_ > X^2^_i_.

In total we identified at least one correct clone for 5661 ORFs (Fig. 1c, Table S1). These clones, one for each ORF, were arrayed in 96-well plates forming the C-SWAT library v1.0.

#### 3. Tag swap with the C-SWAT library

Donor plasmids pMaM482, pMaM484, pYD10, pYD11, pYD13 and pYD14 (Table S3) were transformed into the YMaM639 donor strain.

To test the efficiency of tag swapping (Fig. S2b), we selected 20 C-SWAT strains for highly expressed proteins to facilitate single-cell fluorescence measurements with flow cytometry. The strains were randomly selected from the first plate of the C-SWAT library, which mostly contains C-SWAT strains for highly expressed proteins. Strains with tagged ribosomal subunits, histones or corresponding to proteins expressed at less than 20000 molecules per cell^9^ were excluded. This set of 20 strains was crossed with four donor strains carrying different donor plasmids: YYD2 (type I mNeonGreen donor), YYD3 (type II mNeonGreen donor), YYD8 (type III mNeonGreen donor) and YYD6 (empty type II donor) (Table S4).

The full C-SWAT library was crossed with the following four donor strains: YYD2 (type I mNeonGreen donor), YYD3 (type II mNeonGreen donor), YYD4 (type II mScarlet-I donor) and YYD5 (empty type I donor) (Table S4).

Crossing and subsequent tag swapping were performed by sequentially pinning the strains on appropriate media using a ROTOR HDA pinning robot (Singer Instruments) in 1536-colony format according to the synthetic genetic array (SGA) procedure^10^ as follows:

- mating of C-SWAT and donor strains on YPD plates (10 g/L yeast extract (BD Biosciences, 212750), 20 g/L peptone (BD Biosciences, 211677), 20 g/L glucose (Merck, 108337), 20 g/L agar (BD Biosciences, 214010)), 1 day at 30°C;
- selection of diploids on SC(MSG)-Ura + G-418 plates (1.7 g/L yeast nitrogen base without amino acids and ammonium sulfate (BD Biosciences, 233520), 1 g/L monosodium glutamic acid (MSG) (Sigma-Aldrich, G1626), 2 g/L amino acid mix SC(MSG)-Ura (glutamic acid replaced by MSG), G-418 (200 mg/L, Biochrom, A291-25), 20 g/L glucose, 20 g/L agar), 1 day at 30°C;
- sporulation on SPO plates (20 g/L potassium acetate (Sigma-Aldrich, 25059), 20 g/L agar), 5 days at 23°C;
- selection of haploids, step 1, on SC(MSG)-Ura/His/Arg/Lys + canavanine/thialysine plates (50 mg/L canavanine (Sigma-Aldrich, C1625), 50 mg/L thialysine (Sigma-Aldrich, A2636)), 2 days at 30°C;
- selection of haploids, step 2, on SC(MSG)-Ura/His/Arg/Lys + canavanine/thialysine/G-418 plates (50 mg/L canavanine, 50 mg/L thialysine, 200 mg/L G-418), 2 days at 30°C;
- selection of haploids, step 3, on SC(MSG)-Ura/His/Arg/Lys + canavanine/thialysine/G-418/clonNAT plates (50 mg/L canavanine, 50 mg/L thialysine, 200 mg/L G-418, 100 mg/L clonNAT (Werner BioAgents, 5.0)), 2 days at 30°C;
- induction of tag swapping on SC-His Gal/Raf plates (6.7 g/L yeast nitrogen base without amino acids (BD Biosciences, 291940), 2 g/L amino acid mix SC-His, 20 g/L galactose (Serva, 22020), 20 g/L raffinose (Sigma-Aldrich, R0250), 20 g/L agar), 2 days at 30°C (done twice);
- selection against the acceptor module on SC-His + 5-FOA plates (6.7 g/L yeast nitrogen base without amino acids, 2 g/L amino acid mix SC-His, 20 g/L glucose, 1 g/L 5-FOA (Apollo Scientific, PC4054), 2 days at 30 °C.

Finally, strains resulting from the swap of the full C-SWAT library were pinned on SC-His and grown for 1 day at 30°C prior to fluorescence measurements of colonies.

Strains swapped to test the efficiency of tag swapping were pinned from SC-His + 5-FOA plates either to SC(MSG)-His plates for all three donor types or to SC(MSG)-His + hygromycin plates (200 mg/L Hygromycin B Gold, Invivogen, ant-hg-5) for the type II donor or to SC(MSG)-His + G-418 plates (200 mg/L G-418) for the type III donor. Finally, all strains were pinned on SC-His plates and grown for 1 day at 30°C prior to fluorescence measurements with flow cytometry.

#### 4. Flow cytometry

Strains were grown to saturation in 96-well plates (150 μl of SC-His medium per well) at 30°C, diluted into fresh SC-His medium and grown for 8 h at 30°C to 2-8×10^7^ cells/ml. Fluorescence measurements were performed on a BD FACSCanto RUO (BD Biosciences) equipped with a high-throughput sampler loader, a 488 nm laser and a combination of 505 nm long-pass and 530/30 nm band pass emission filters for mNeonGreen detection. Populations were gated for single cells in the G1 phase of the cell cycle using the first peak in the side scatter width (SSC-W) histogram and 20000 cells were measured for each strain.

#### 5. Colony fluorescence measurements

Strains resulting from the swap of the full C-SWAT library with mNeonGreen and mScarlet-I donors in three technical replicates were arranged next to each other and three technical replicates of a negative control (tag swap using the empty donor plasmid pMaM482). Strains were pinned on SC-His agar plates using a ROTOR HDA pinning robot (Singer Instruments) and grown at 30°C for 24 h. Fluorescence measurements were performed at 30°C with Infinite M1000 or Infinite M1000 Pro plate readers (Tecan) equipped with stackers for automated plate loading and custom temperature control chambers. Detector gain was set manually to avoid saturation and measurements were performed at 400 Hz frequency of the flash lamp, with ten flashes averaged for each measurement, in two channels: mScarlet-I (569/10 nm excitation, 593/10 nm emission) and mNeonGreen (506/5 nm excitation, 517/5 nm emission). Measurements were filtered for potentially failed crosses based on colony size after haploid selection. Colony area measurements for each individual plate were median-centered prior to calculation of median colony size for the entire data set. Scaled median absolute deviation (MAD) served as a robust estimate of standard deviation, and colonies within the 0.5th percentile of a normal distribution centered at the median with a spread of scaled MAD were defined as failed crosses. Tag swaps with less than two successfully crossed replicates were removed from the analysis. Fluorescence intensities for each plate were normalized to the median fluorescence of a reference strain set that was present on every plate. Intensities of sample colonies were either corrected for background by subtracting the average intensity of negative control colonies or expressed in background units, i.e. divided by the average intensity of negative control colonies (Table S2a).

#### 6. Fluorescence microscopy

Strains were inoculated in 96-well plates in synthetic complete (SC) low-fluorescence medium (SC medium prepared with yeast nitrogen base lacking folic acid and riboflavin^11^) and grown at 30°C to mid log phase for 7-8 h. 150 μl of each culture were used for microscopy in glass-bottom 96-well plates (MGB096-1-2-LG-L; Matrical) coated with concanavalin A, as described^12^. Imaging was performed on a Nikon Ti-E widefield epifluorescence microscope with a 60x ApoTIRF oil immersion objective (1.49 NA, Nikon), an LED light engine (SpectraX, Lumencor), an sCMOS camera (Flash4, Hamamatsu) and an autofocus system (Perfect Focus System, Nikon) with either bright field, 469/35 excitation and 525/50 emission filters or 542/27 excitation and 600/52 emission filters (all from Semrock except 525/50, which was from Chroma). Z-stacks of 11 planes with 0.5 μm spacing were recorded with two different exposure times for mNeonGreen and mScarlet-I.

#### 7. Availability of resources

The C-SWAT library, the derived mNG-I, mNG-II and mSC-II libraries, all reagents necessary to use the C-SWAT library for high-throughput strain construction and custom scripts for analysis of Anchor-Seq data are available upon request.

## Supplementary Figures

**Figure S1.**
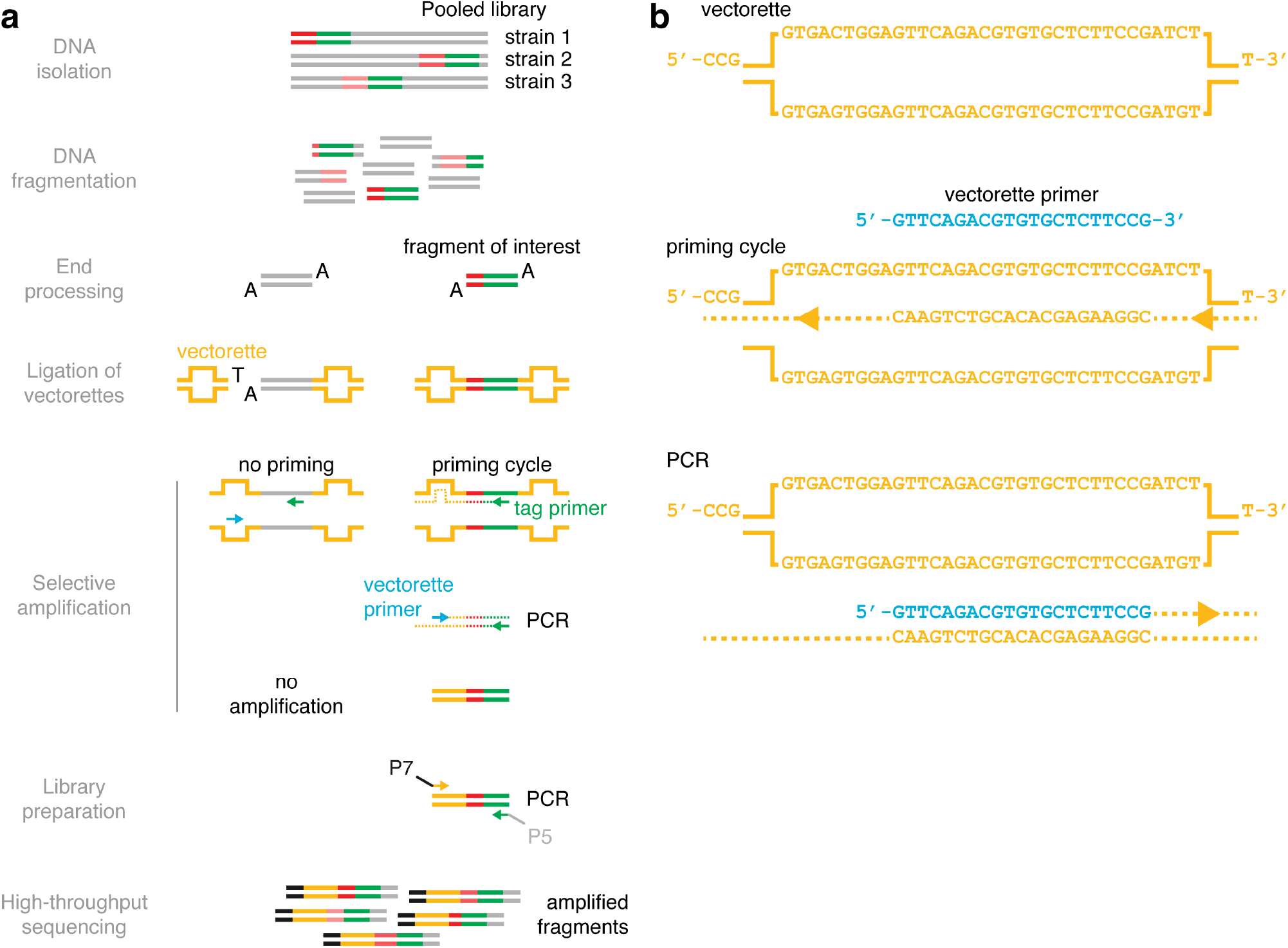
Anchor-Seq workflow. **(a)** Genomic DNA is extracted from a pooled library, where a different ORF is modified (*e.g*., tagged with the C-SWAT acceptor module) in each strain. The DNA is sheared and the fragment ends are processed to allow A-T ligation of vectorette adaptors. Selective amplification of fragments containing the library tag is achieved with two primers: a tag primer (green) that anneals to the tag sequence and a vectorette primer (red) that has the same sequence as the vectorette adaptor and thus cannot anneal to it. The sequence complementary to the vectorette primer is produced in the first PCR cycle initiated by the tag primer. Exponential amplification only of fragments containing the library tag can subsequently occur (15 PCR cycles in total). The PCR product is purified and subjected to another 15 PCR cycles to add the P5 and P7 Illumina sequences to the amplified fragments. **(b)** Sequence of the vectorette primer (blue) in relation to the central portion of the vectorette adaptor and the product of the priming cycle with the tag primer (selective amplification step in (a)).

**Figure S2.**
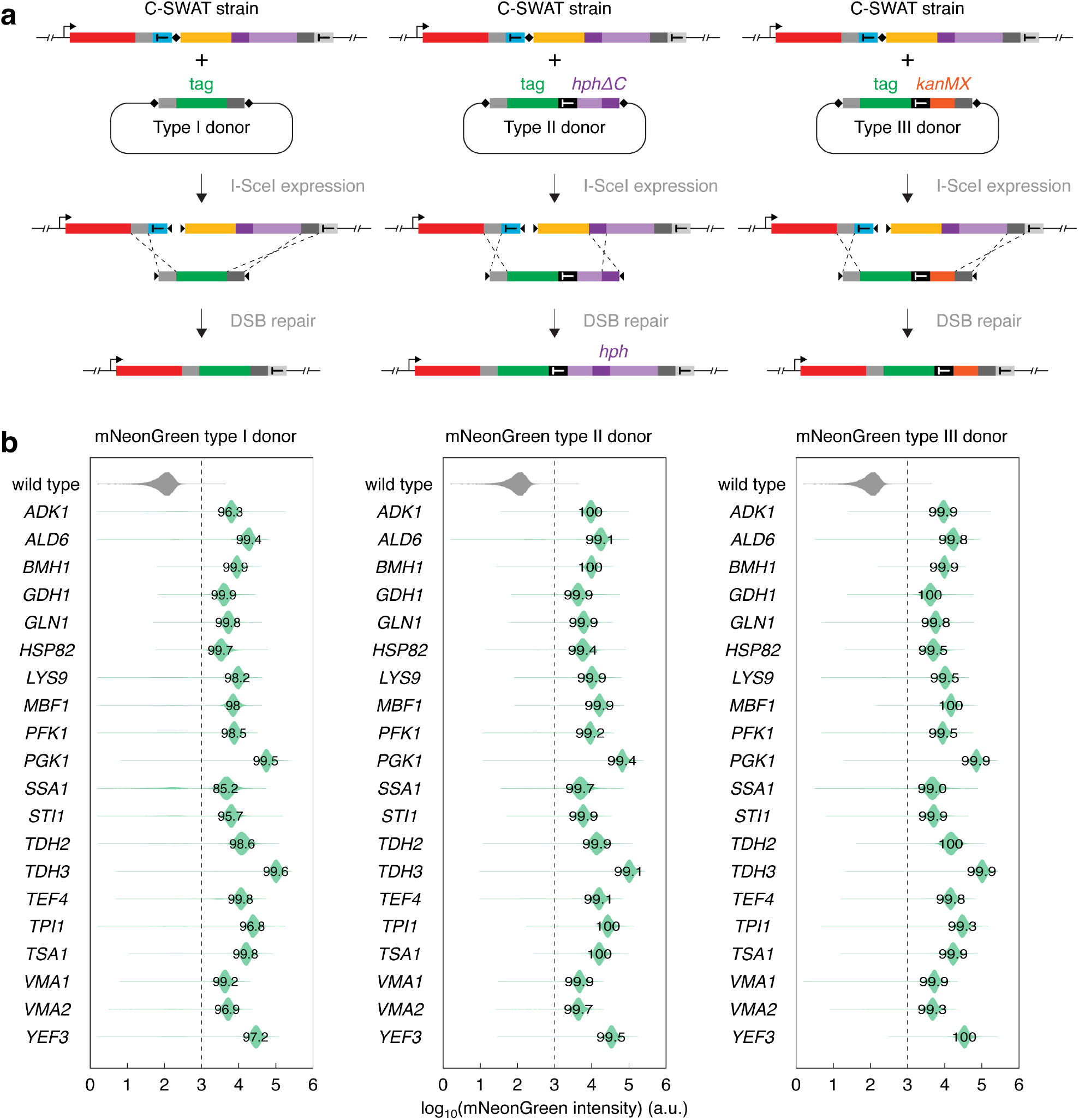
Efficiency of tag swapping with different donors. **(a)** Three types of donor plasmids for tag swapping with C-SWAT strains. The type I donor is used for seamless tagging. In this donor the tag is placed between short homology arms (gray) and flanked by I-SceI cut sites (black squares). Tag swapping events are selected by resistance to 5-fluoroorotic acid (5-FOA), which indicates loss of the *URA3* selection marker. The type II donor is used for non-seamless tagging. In this donor the left homology arm (light gray) is followed by the tag, the terminator sequence of the *ADH1* gene from *Saccharomyces cerevisiae*, the promoter sequence of the *TEF* gene from *Ashbya gossypii* and the *hphΔC* sequence coding for an N-terminal fragment (amino acids 1-192) of the *hph* (hygromycin resistance-encoding gene) marker. Tag swapping events are selected by resistance to 5-FOA and later/alternatively hygromycin, which indicates reconstitution of the *hph* selection marker. The type III donor is used for non-seamless tagging. In this donor the left homology arm (light gray) is followed by the tag, the *ADH1* terminator sequence and a selection marker (*e.g*., the *kanMX* marker (resistance to G-418)). Tag swapping events are selected by resistance to 5-FOA and later G-418. **(b)** Comparison of tagging efficiency using three donors with the mNeonGreen fluorescent protein. Protein tagging was performed with 20 C-SWAT strains for the indicated genes. Distributions of single-cell mNeonGreen fluorescence intensities measured with flow cytometry (~20000 cells per strain). The percentage of cells with fluorescence above background (fluorescence of a wild type strain, dashed line) is indicated in the plots.

**Figure S3.**
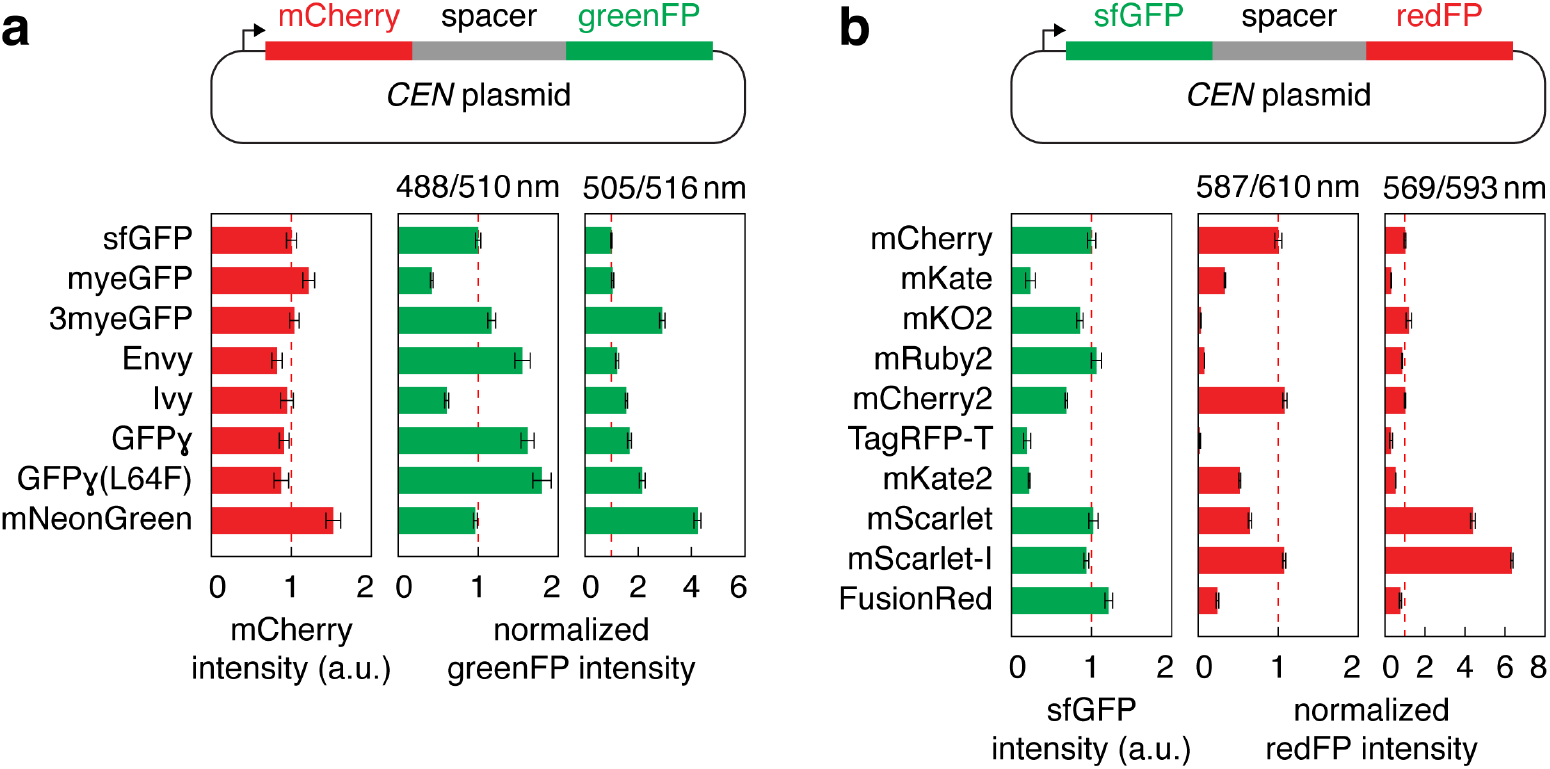
Brightness of different fluorescent proteins in yeast. **(a)** Relative brightness of different green fluorescent proteins (greenFPs) in yeast. Each greenFP was fused to mCherry, with Don1 as a spacer protein to minimize Förster resonance energy transfer (FRET) between the two fluorescent proteins. The fusions were expressed in yeast from the strong constitutive GPD promoter. Whole colony greenFP fluorescence intensities, measured with two sets of excitation and emission wavelengths, were normalized for protein expression levels using mCherry fluorescence intensities. The resulting relative brightness estimates were normalized to sfGFP (mean ± s.d., n = 5 biological replicates each with 4 technical replicates). Based on these results and excitation/emission spectra, we estimate that mNeonGreen with 505/516 nm excitation/emission is ~2 fold brighter than sfGFP with 488/510 nm excitation/emission. **(b)** Relative brightness of different red fluorescent proteins (redFPs) in yeast. Each redFP was fused to sfGFP with Don1 as a spacer protein. The fusions were expressed from the GPD promoter and whole colony redFP fluorescence intensities, measured with two sets of excitation and emission wavelengths, were normalized for protein expression levels using sfGFP fluorescence intensities. The resulting relative brightness estimates were normalized to mCherry (mean ± s.d., n = 3 biological replicates each with 4 technical replicates). Based on these results and excitation/emission spectra, we estimate that mScarlet-I with 569/593 nm excitation/emission is ~3 fold brighter than mCherry with 587/610 nm excitation/emission.

**Figure S4.**
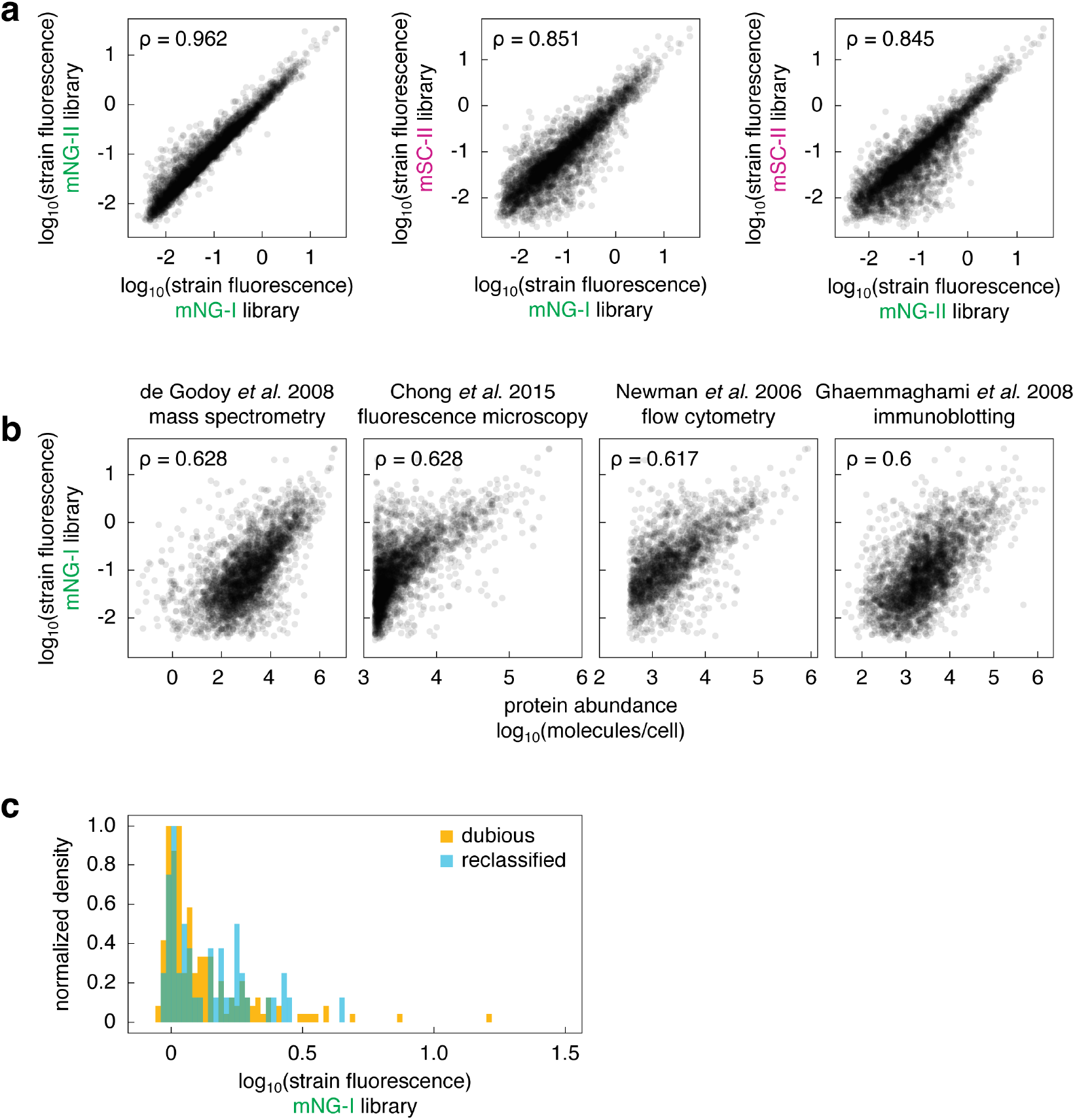
Measurements of protein abundance with mNeonGreen and mScarlet-I libraries. **(a)** Comparison of relative protein expression levels measured with the mNG-I, mNG-II and mSC-II libraries. The libraries were derived from the C-SWAT library by tag swapping with the three donor plasmids in Fig. 2a. Fluorescence intensities of colonies were corrected for background fluorescence. **(b)** Comparison between relative protein expression levels measured with the mNG-I library and estimates of absolute protein abundance from ref. 9. Absolute estimates are based on measurements of protein abundance with mass spectrometry of wild type yeast^13^, fluorescence microscopy and flow cytometry of strains expressing GFP fusions^14,15^ or immunoblotting of TAP-tagged proteins^16^. Only strains with fluorescence intensities at least 1.2 fold above background were used in (a) and (b). **(c)** Fluorescence levels of mNG-I strains for two groups of ORFs: annotated as *dubious* as of January 2017 (dubious) and reclassified from *dubious* to *verified* or *uncharacterized* between July 2016 and January 2017 (reclassified). Fluorescence intensities of colonies are expressed in units of background fluorescence, not corrected for background fluorescence.

## Supplementary Tables

**Table S1**. C-SWAT library description. S2/S3 primers and number of positive clones validated by Anchor-Seq for each ORF.

**Table S2**. Protein expression levels in mNG-I, mNG-II and mSC-II libraries. **(a)** Fluorescence intensities of colonies, fold over background or background-corrected. **(b)** Expression and localization of previously undetected ORFs. **(c)** Changes in classification of dubious ORFs.

**Table S3.**
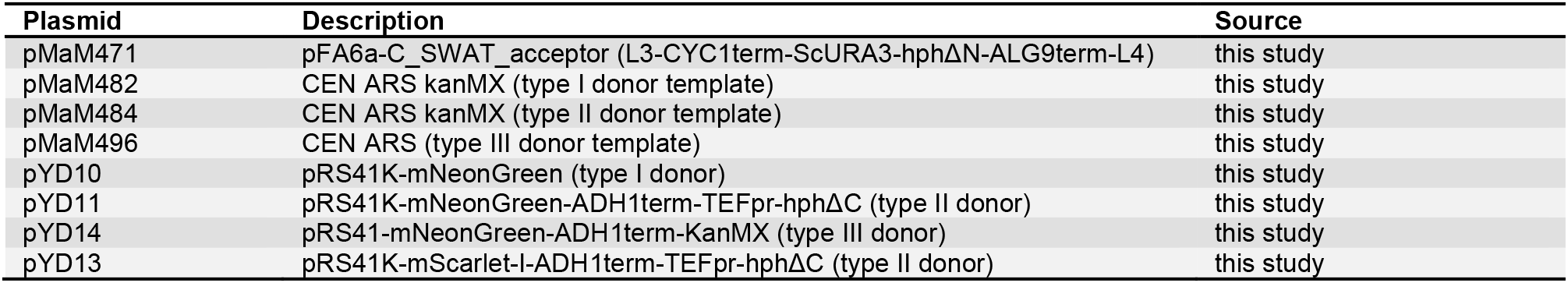
Plasmids.

| Plasmid | Description | Source |
| --- | --- | --- |
| pMaM471 | pFA6a-C_SWAT_acceptor (L3-CYC1term-ScURA3-hphΔN-ALG9term-L4) | this study |
| pMaM482 | CEN ARS kanMX (type I donor template) | this study |
| pMaM484 | CEN ARS kanMX (type II donor template) | this study |
| pMaM496 | CEN ARS (type III donor template) | this study |
| pYD10 | pRS41K-mNeonGreen (type I donor) | this study |
| pYD11 | pRS41K-mNeonGreen-ADH1term-TEFpr-hphΔC (type II donor) | this study |
| pYD14 | pRS41-mNeonGreen-ADH1term-KanMX (type III donor) | this study |
| pYD13 | pRS41K-mScarlet-I-ADH1term-TEFpr-hphΔC (type II donor) | this study |

**Table S4.** Yeast strains.

| Strain | Background | Genotype | Source |
| --- | --- | --- | --- |
| BY4741 | S288c | *MATa his3Δ1 leu2Δ0 met15Δ0 ura3Δ0* | ref. 1 |
| Y8205 | S288c | *MATalpha can1Δ::STE2pr-SpHIS5 and lyp1Δ::STE3pr-LEU2 his3Δ1 leu2Δ0 ura3Δ0* | ref. 4 |
| YMaM639 | Y8205 | *leu2Δ0::GAL1pr-NLS-I-SceI-natNT2* | this study |
| YYD2 | YMaM639 | pYD10 (type I mNeonGreen donor) | this study |
| YYD3 | YMaM639 | pYD11 (type II mNeonGreen donor) | this study |
| YYD8 | YMaM639 | pYD14 (type III mNeonGreen donor) | this study |
| YYD4 | YMaM639 | pYD13 (type II mScarlet-I donor) | this study |
| YYD5 | YMaM639 | pMaM482 (empty type I donor) | this study |
| YYD6 | YMaM639 | pMaM484 (empty type II donor) | this study |

**Table S5.**
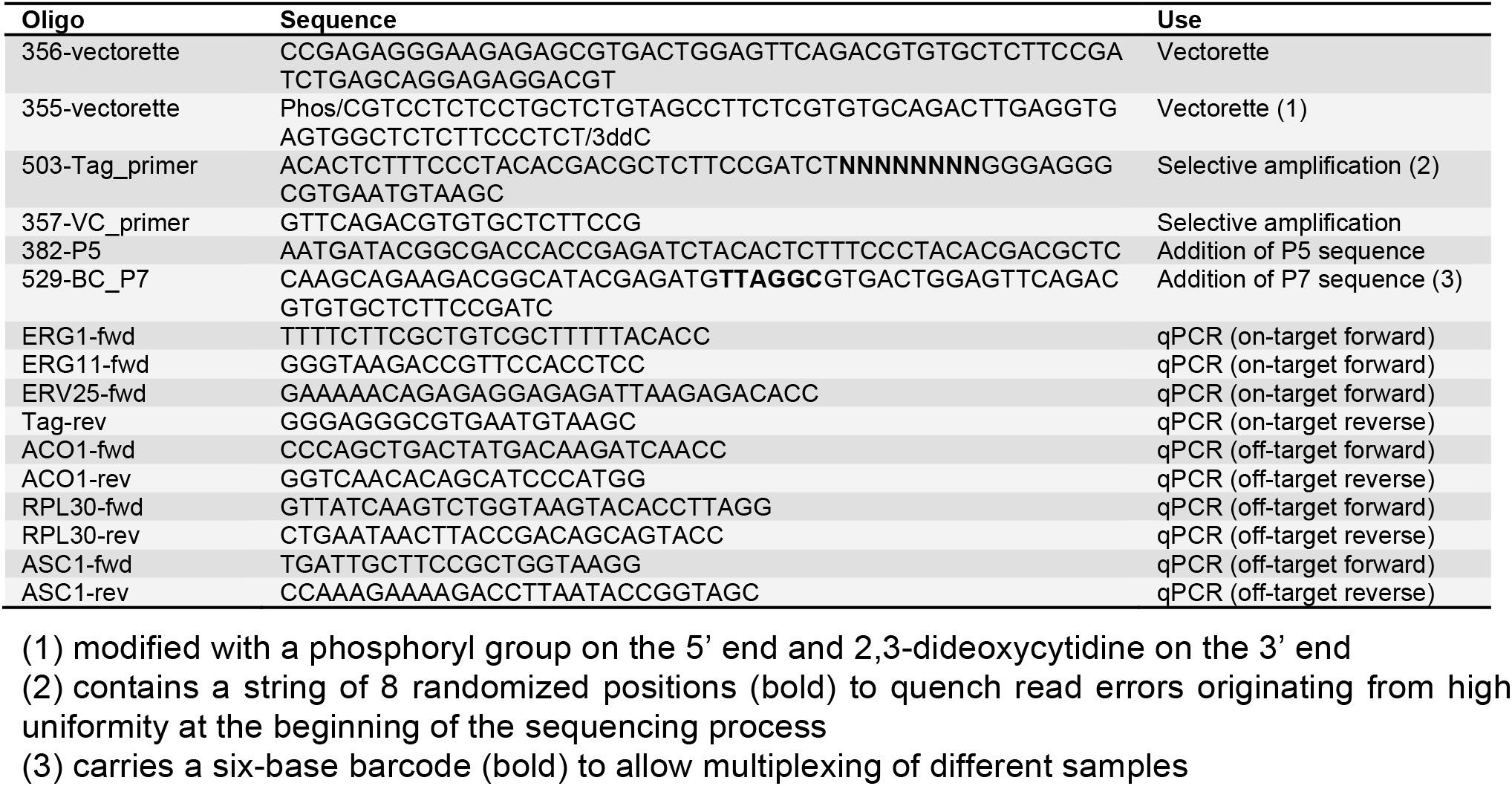
Oligonucleotides.

